# Seasonal influenza viruses decay more rapidly at intermediate humidity in droplets containing saliva compared to respiratory mucus

**DOI:** 10.1101/2023.07.11.548566

**Authors:** Nicole C. Rockey, Valerie Le Sage, Linsey C. Marr, Seema S. Lakdawala

## Abstract

Expulsions of virus-laden aerosols or droplets are an important source of onward respiratory virus transmission and can originate from both the oral and nasal cavity of an infected host. However, the presence of infectious influenza virus in the oral cavity during infection has not been widely considered, and thus little work has explored the environmental persistence of influenza virus in oral cavity expulsions that may facilitate transmission. Using the ferret model, we detected infectious virus in the nasal and oral cavities, suggesting that virus can be expelled into the environment from either anatomical site. We also assessed the stability of two influenza A viruses (H1N1 and H3N2) in droplets of human saliva or respiratory mucus over a range of relative humidities. We observed that influenza virus infectivity decays rapidly in saliva droplets at intermediate relative humidity, while viruses in airway surface liquid droplets retain infectivity. Virus inactivation was not associated with bulk protein content, salt content, or droplet drying time. Instead, we found that saliva droplets exhibited distinct inactivation kinetics during the wet and dry phases at intermediate relative humidity and that droplet residue morphology may lead to the elevated first-order inactivation rate observed during the dry phase. Additionally, distinct differences in crystalline structure and nanobead localization were observed between saliva and airway surface liquid droplets. Together, our work demonstrates that different respiratory fluids exhibit unique virus persistence profiles and suggests that influenza viruses expelled from the oral cavity may contribute to virus transmission in low and high humidity environments.

**Importance:** Determining how viruses persist in the environment is important for mitigating transmission risk. Expelled infectious droplets and aerosols are composed of respiratory fluids, including saliva and complex mucus mixtures, but how influenza viruses survive in such fluids is largely unknown. Here, we find that infectious influenza virus is present in the oral cavity of infected ferrets, suggesting that saliva-containing expulsions can play a role in onward transmission. Additionally, influenza virus in droplets composed of saliva degrades more rapidly than virus within respiratory mucus. Droplet composition impacts the crystalline structure and virus localization in dried droplets. These results suggest that viruses from distinct sites in the respiratory tract could have variable persistence in the environment, which will impact viral transmission fitness.

## Introduction

Influenza virus presents a substantial global disease burden, having caused most of the major pandemics in the last century and being responsible for seasonal outbreaks that result in significant morbidity and mortality (1). Understanding how influenza viruses can transmit efficiently between hosts is therefore critical to better mitigating the circulation of these viruses in future pandemics and ongoing seasonal epidemics.

Influenza virus is spread through the expulsion of virus-containing respiratory fluid aerosols or droplets that remain infectious in the environment and reach the respiratory tract of a new host to initiate a new round of viral replication (2, 3). Respiratory activities, such as breathing, coughing, talking, and sneezing, all produce aerosols or droplets spanning a wide range of sizes that may contain infectious virus (4, 5). Many studies have indeed confirmed the presence of influenza virus RNA or infectious influenza virus in aerosols expelled from infected individuals (6–10). These virus-containing aerosols or droplets are comprised of respiratory fluids originating from distinct parts of the respiratory tract, including the lungs, trachea, oral cavity, and nasal cavity (4). Importantly, the relative levels of infectious virus emitted from different parts of the respiratory tract and their environmental persistence in distinct respiratory fluids are not well understood.

To transmit efficiently, expelled viruses must retain infectivity in the environment. The impact of certain environmental factors, such as humidity and temperature, on virus persistence in droplets or aerosols has been studied extensively (11–18). Work to date has demonstrated an overall reduction in persistence of enveloped viruses at intermediate humidities and ambient temperatures in both aerosols and droplets (13, 14, 16–18), although the inverse phenomenon has been observed in some work with nonenveloped viruses (11, 18). Unfortunately, these studies have primarily been conducted using laboratory-derived solutions (e.g., cell culture medium, phosphate buffered saline solution), limiting the relevance of these findings in informing real-world transmission scenarios.

Some studies have recently begun to employ physiologically relevant aerosol or droplet matrices. Influenza virus has been found to maintain infectivity in aerosols and droplets comprised of human airway surface liquid collected from human bronchial epithelial cell cultures differentiated at an air-liquid interface, representative of respiratory fluid from the lower respiratory tract, at relative humidities (RH) ranging from ∼ 20% to over 90% (19, 20). Similarly, a recent publication using nebulized saliva microdroplets containing human virus surrogates, including bacteriophages MS2, phiX174, and phi6, observed an overall increase in persistence of virus in saliva droplets at all RHs compared to water or media after 14 hours (21). However, studies of other mammalian viruses in saliva have found that viruses such as murine coronavirus and vesicular stomatitis virus decay rapidly at intermediate RHs (22, 23). More research is clearly required to understand how different respiratory viruses retain infectivity in distinct respiratory fluid matrices under a variety of environmental conditions.

The differences in environmental virus inactivation exhibited across different RHs, viruses, and compositions may be due to various factors, including salt or protein content, evaporation kinetics, and/or pH. Prior research has focused on the influence of solutes and proteins on environmental virus decay (11, 15), suggesting that protein may provide a protective effect at intermediate relative humidity, while salt may inactivate viruses during the wet phase of droplet drying (15). The role of pH in virus inactivation has been central to recent virus persistence studies, although findings are not consistent. Modeling work demonstrated that rapid acidification of aerosols that can render viruses such as SARS-CoV-2 or influenza virus inactive, while another publication showed SARS-CoV-2 inactivation correlated with an increase in pH at high RH (24, 25). Experiments in saliva have suggested model viruses are protected by carbohydrates at low RH (22) or susceptible to antiviral proteins at intermediate RH (23). Despite these efforts, the mechanisms driving virus inactivation in distinct respiratory fluid fluids remain largely unknown and may vary depending on virus type. A better understanding of the factors governing viral persistence is important to develop more informed interventions that limit respiratory virus transmission.

In this study, we first established whether influenza virus is present in the oral cavity of experimentally infected ferrets and found substantial levels of infectious virus present in saliva. Given the potential for salivary particles to mediate transmission, we examined the impact of saliva on the environmental stability of influenza virus. To this end, we assessed the persistence of two influenza A viruses, an H3N2 virus (A/Perth/16/2009 (H3N2); H3N2) and the H1N1 2009 pandemic virus (A/California/07/2009 (H1N1); H1N1pmd09), in 1 μL droplets comprised of human saliva or airway surface liquid at low, medium, and high RH (i.e., 20%, 50%, and 80%) and ambient temperature over a two-hour period. We observed distinct virus decay and droplet morphology patterns in saliva compared to airway surface liquid. Our findings demonstrate the importance of using physiologically relevant matrices when assessing environmental persistence and lend insights into the mechanisms driving virus persistence in different respiratory fluids. Taken together, this information can ultimately be used to help inform strategies for restricting the spread of influenza viruses.

## Results

### Oral swabs from infected ferrets contain infectious influenza virus

The concentrations of infectious influenza virus found in the saliva of infected individuals over time has not been widely characterized. We therefore assessed the infectious virus levels in the oral cavity of ferrets infected intranasally with two seasonal influenza viruses, H1N1pdm09 and H3N2. Specifically, we swabbed each ferret’s tongue, cheeks, hard palate, and soft palate; importantly, we did not swab the back of the throat, which could include virus from the lower respiratory tract. In parallel, we sampled the ferret nasal cavity by nasal wash to observe the relative amount of virus present in the proximal tip of the ferret nostril.

Influenza virus concentrations from oral swabs followed the same trends observed in nasal wash levels over the course of infection for both H1N1pdm09 and H3N2 viruses, although infectious virus in oral swabs fell below the limit of detection earlier in infection for H3N2 infection (Fig. 1). Infectious influenza virus was consistently detected in oral swabs from intranasally infected ferrets on days one through five post-infection, with levels as high as ∼ 4-log_10_ TCID_50_/swab. Viral titers in the nasal wash of some ferrets were as high as 5.5-log_10_ TCID_50_/mL on days one or two post-infection. Together, these results demonstrate that infectious influenza virus is present at elevated levels in saliva within the oral cavity during infection. Expulsion of influenza viruses in saliva droplets or aerosols during breathing, talking, or coughing may therefore act as a source of onward transmission.

**Figure 1.**
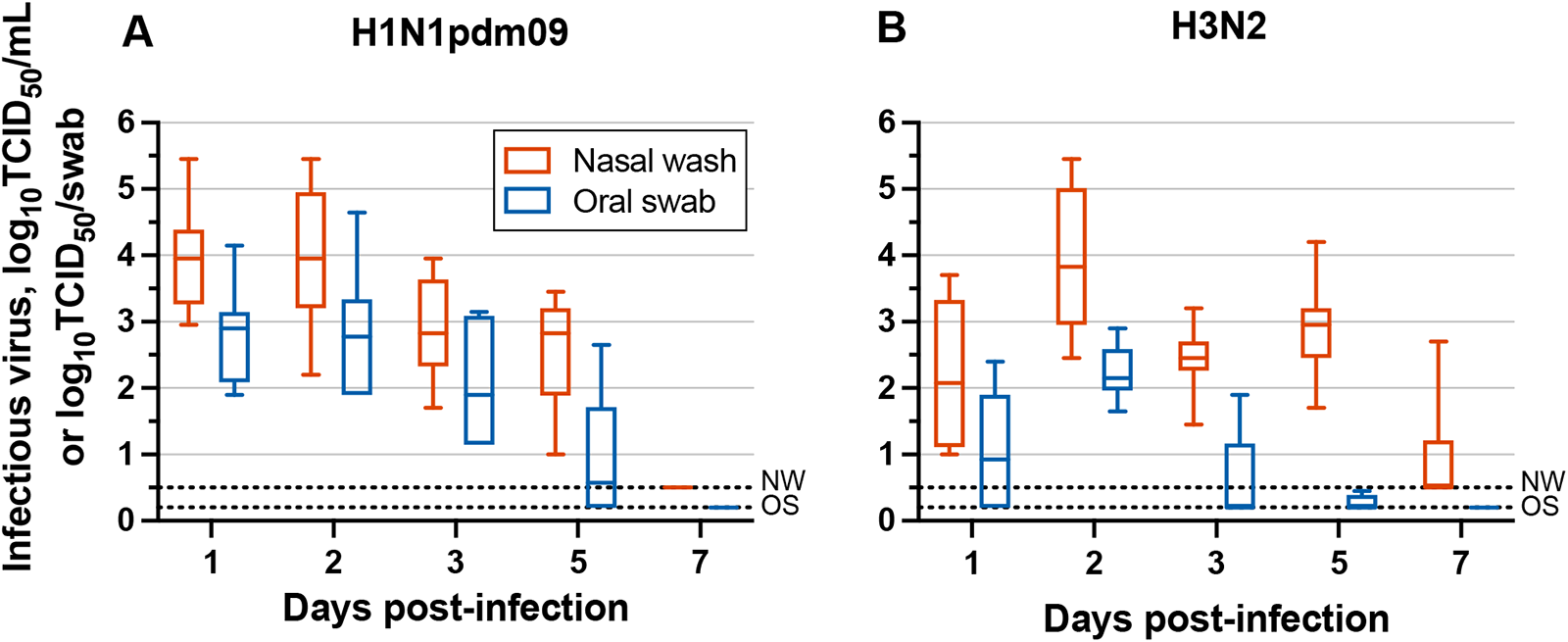
Infectious influenza A virus levels in the nasal (red) and oral (blue) cavities of ferrets following infection. Ferrets were infected via intranasal inoculation with (A) H1N1pdm09 or (B) H3N2. Individual replicates (n = 8) are included for each box and whisker plot. Dashed lines represent the limit of detection for nasal wash samples (NW) and oral swab samples (OS).

### Influenza A viruses decay rapidly at intermediate relative humidity in saliva droplets but not in airway surface liquid droplets

The presence of influenza virus in the host oral cavity on multiple days in experimentally infected animals highlights the relevance of saliva in transmission. However, the persistence of influenza virus in saliva has not been characterized. We therefore investigated the environmental stability of two influenza A virus strains, H1N1pdm09 and H3N2, in 10 x 1 μL saliva droplets after one or two hours of exposure to 20%, 50%, and 80% RH and ambient temperature (i.e., ∼ 22°C) (Fig. 2 and Fig. S1). Virus decay in saliva was compared to decay in airway surface liquid, which serves as a biological surrogate for respiratory airway secretions. Our group has previously demonstrated reduced decay of human seasonal influenza viruses in droplets and aerosols consisting of airway surface liquid as compared to cell culture medium (19, 20).

Major trends in virus inactivation were similar for both influenza A virus strains evaluated (Fig. 2). As expected, influenza viruses in airway surface liquid droplets were protected from decay, never exceeding 1.1-log_10_ decay, on average, over two hours. We found that influenza A virus persistence was greatest at 20% RH for both droplet compositions, with maximum average decay for both strains reaching just 1-log_10_ after two hours (Fig. 2A and 2D). Interestingly, we observed increased decay of influenza A virus in saliva droplets containing saliva at 50% and 80% RH compared to airway surface liquid droplets (Fig. 2B – C; Fig. 2E – F). At one hour of exposure to 50% RH, decay of infectious H1N1pdm09 and H3N2 in saliva droplets was significantly greater than in airway surface liquid (Fig. 2B and 2E). At two hours, virus decay differences for both strains at 50% RH varied significantly by droplet composition, with average influenza virus degradation in saliva droplets exceeding 3-log_10_, while virus decay in airway surface liquid droplets was only ∼ 1-log_10_. Virus decay in saliva droplets at 80% RH was intermediate, with average decay of 2.9-log_10_ and 1.7-log_10_ for H1N1pdm09 and H3N2 viruses, respectively (Fig. 2C and 2F). Our findings establish that influenza A virus persistence in droplets is highly composition dependent. In addition, these two influenza A subtypes decay in a similar manner for a given RH and droplet composition.

**Figure 2.**
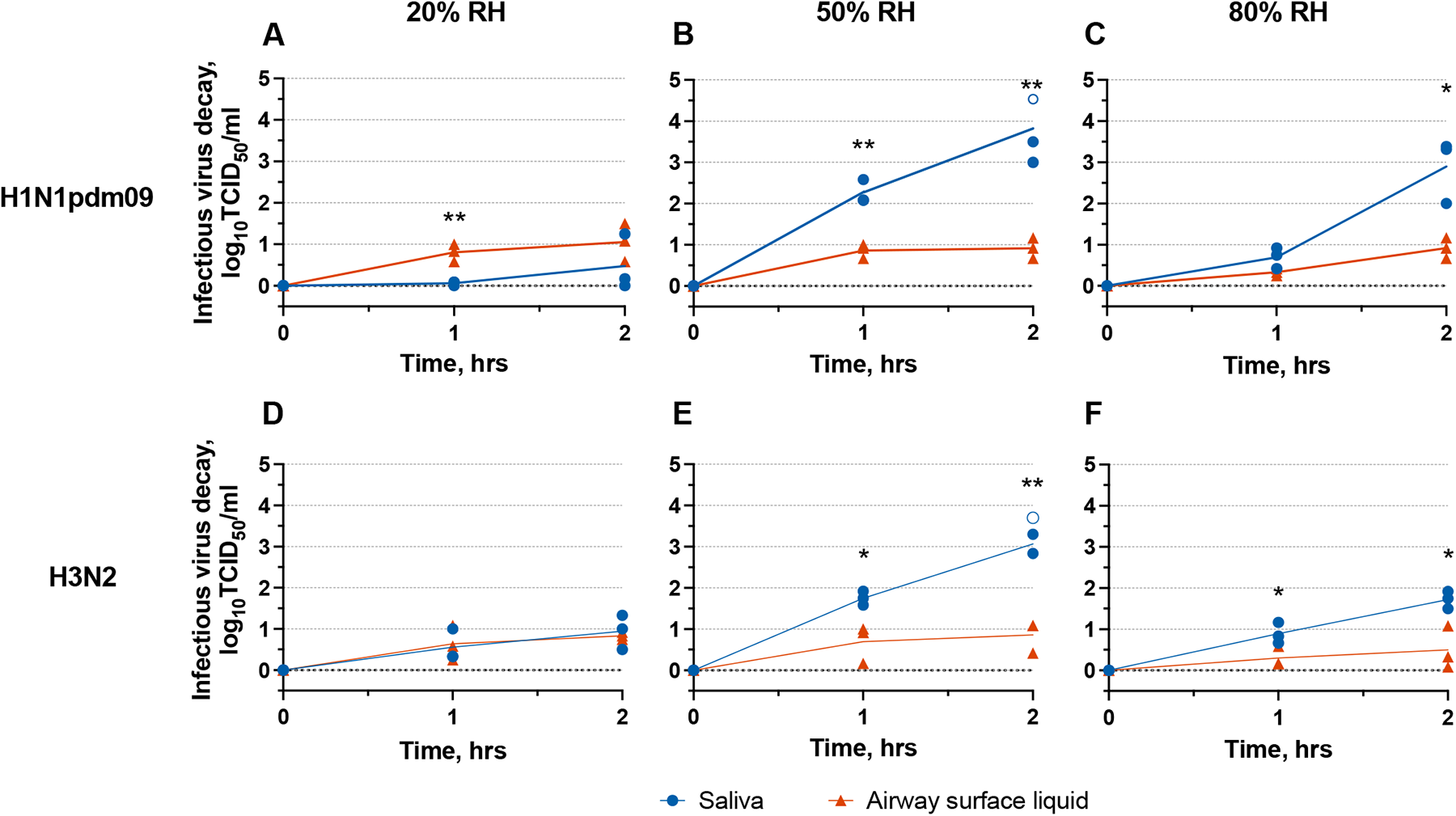
Reductions in infectious influenza A virus in droplets comprised of saliva or airway surface liquid. H1N1pdm09 or H3N2 log_10_ decay at 20%, 50%, or 80% RH after one and two hours of exposure. Open circles indicate infectious virus decay beyond detection limits. Three technical replicates were conducted for three independent replicates, and the mean decay of three technical replicates is shown for each independent replicate. Unpaired t-tests were conducted to establish when influenza virus decay was significantly different between respiratory fluid droplets. * = p < 0.05; ** = p < 0.005.

To ensure the observed decay was not a side effect of poor virus recovery, we measured RNA recovered from virus-laden droplets at 50% RH in saliva and airway surface liquid at zero and one hour. Results demonstrate no significant loss in viral RNA levels over this period (Fig. S2A; p > 0.05), suggesting that any loss of infectivity observed in our experiments is indeed a result of virus inactivation in droplets (Fig. S2B). Because gene copy concentrations were similar over the droplet drying period, but infectious virus levels were reduced, the gene copy to infectious unit ratio increased with time (Fig. S2C). This is an important consideration when using genome-based detection methods to study virus persistence, because genome copy levels are likely to overestimate infectious virus levels following longer periods of environmental exposure.

### Influenza virus inactivation is not driven by phase, salt content, or protein concentration of the droplet

The inactivation mechanisms driving the differences in virus persistence observed for distinct respiratory fluid droplets are not known. Protein concentration has been suggested to impact virus decay in droplets (15); however, the airway surface liquid and human saliva used in our current study have similar levels of total protein content (Table 1). Therefore, differences in virus decay by droplet composition are not due to overall protein levels, although specific proteins within saliva or airway surface liquid could be responsible. In addition, protein partitioning (e.g., to the air-liquid interface or phase interphases within a droplet) or aggregation could differ between saliva and airway surface liquid, which could also contribute to the observed differences in persistence. Conductivity, a measure of dissolved ions, was also assessed to establish the solute concentrations in these respiratory fluids. Human saliva had lower conductivity, 3.54 mS/cm, than airway surface liquid, with a level of 16.52 mS/cm (Table 1). Elevated salt (i.e., ion) concentrations have been linked to increased virus inactivation (16). Here, we observed the inverse relation, suggesting that other factors were more important in driving virus inactivation in saliva droplets at 50% RH.

Recent work has shown that acidification of droplet or aerosol pH significantly impacts virus persistence (24, 25). All bulk solutions used in our droplet persistence experiments had a pH that was neutral to slightly basic (Table 1), suggesting that pH levels may not be responsible for the observed decay. Due to technical limitations, we did not measure pH in 1 μL droplets during drying, so we do not know how the pH might have changed during that time. Additionally, the pH in small droplets may differ from that of the bulk solution (26), so we cannot discern how pH might have affected virus inactivation in these experiments.

**Table 1.**
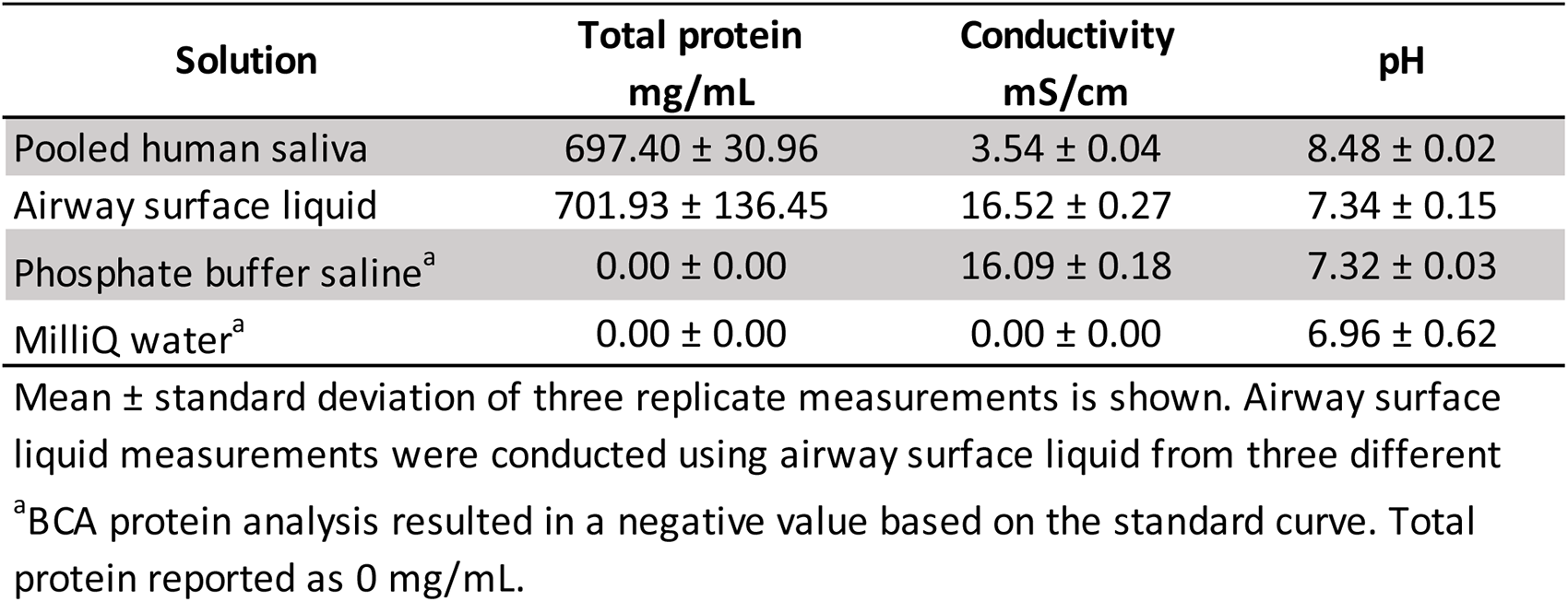
Bulk properties of distinct respiratory fluids, phosphate buffer, and milliQ water.

| Solution | Total protein<br>mg/mL | Conductivity<br>mS/cm | pH |
| --- | --- | --- | --- |
| Pooled human saliva | 697.40 ± 30.96 | 3.54 ± 0.04 | 8.48 ± 0.02 |
| Airway surface liquid | 701.93 ± 136.45 | 16.52 ± 0.27 | 7.34 ± 0.15 |
| Phosphate buffer saline<sup>a</sup> | 0.00 ± 0.00 | 16.09 ± 0.18 | 7.32 ± 0.03 |
| MilliQ water<sup>a</sup> | 0.00 ± 0.00 | 0.00 ± 0.00 | 6.96 ± 0.62 |

Mean ± standard deviation of three replicate measurements is shown. Airway surface liquid measurements were conducted using airway surface liquid from three different <sup>a</sup>BCA protein analysis resulted in a negative value based on the standard curve. Total protein reported as 0 mg/mL.

Beyond salt and protein content of droplets, previous work has suggested virus inactivation in droplets differs depending on whether the droplet is in the wet phase, when it is still evaporating, or the dry phase, when the droplet is no longer losing water (13, 27). We therefore assessed the drying time by measuring the mass of 10 x 1 µL droplets of saliva and airway surface liquid containing H1N1pdm09 over time (Fig. 3) (13, 27). We also ascertained the drying time of the droplets visually by identifying the time at which droplets began to develop a visible white film (SI Videos S1 – S6).

**Figure 3.**
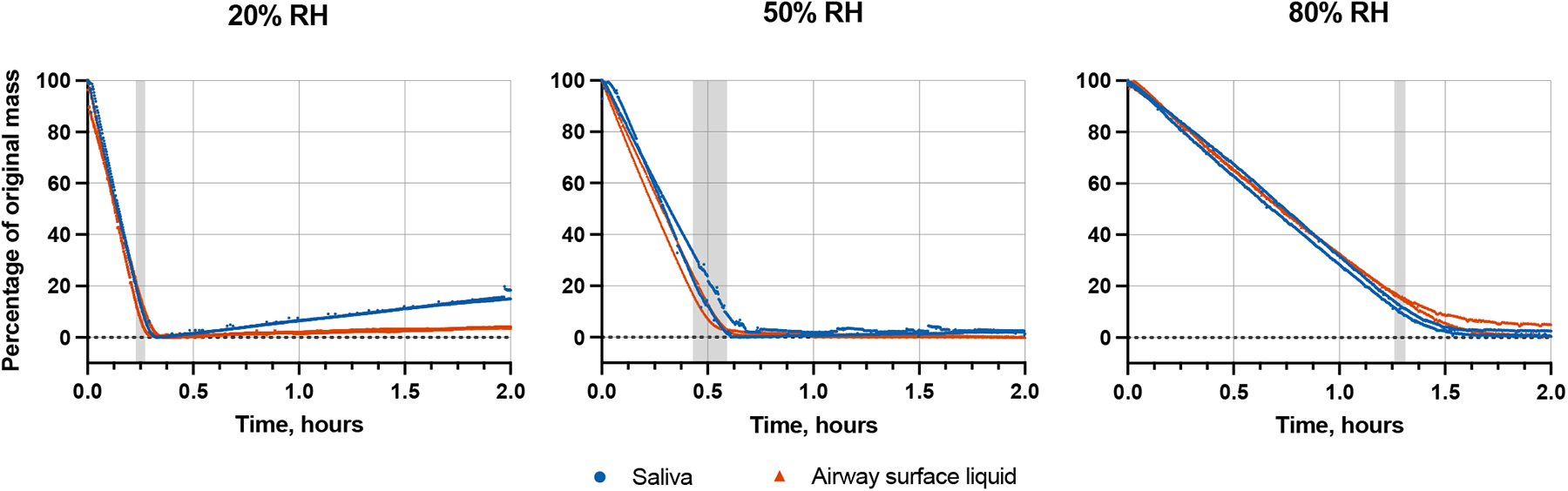
Change in droplet mass over time. Saliva or airway surface liquid droplets containing H1N1pdm09 were exposed to 20%, 50%, or 80% RH for two hours. Shaded regions designate the drying time. Two independent replicates are shown for each RH condition and droplet composition.

As expected, the drying time increased with rising RH. Evaporation kinetics were similar in saliva and airway surface liquid droplets at all tested RHs (Fig. 3, Table S1). Specifically, average drying times differed by at most 4.2 minutes between airway surface liquid and saliva droplets for a given RH. Average drying time at 20% RH in airway surface liquid and saliva was 15 mins and 15.6 mins, respectively, while drying time at 50% RH was slightly greater, 28.2 mins for airway surface liquid and 32.4 mins for saliva. In some trials at 20% RH, droplet mass reached a minimum and then gradually increased. We have observed this behavior before (27) and believe it is due to uptake of water vapor by hygroscopic salts that are exposed upon drying. At 80% RH, droplet mass stabilized after ∼ 1.3 hours in both saliva and airway surface liquid. The drying times defined by mass and visual inspection were similar. The only exception was at 80% RH, where droplets did not dry by observation within the two-hour exposure period. As drying time across droplet matrices was similar (Fig. 3) while decay rates differed (Fig. 2), we can conclude that virus inactivation is not solely a function of evaporation kinetics and that other mechanisms are important.

### Droplet composition influences influenza virus inactivation kinetics before and after drying

To better understand influenza virus inactivation kinetics during the wet and dry phases, we examined the short-term inactivation kinetics of H1N1pdm09 in droplets at 50% RH while drying. Decay of influenza virus in saliva droplets after one hour at 50% RH was ∼ 2-log_10_, while virus in airway surface liquid droplets exhibited significantly less overall decay, ∼ 0.6-log_10_ on average (Fig. 4). Trends in saliva droplet virus decay were distinct across the wet and dry phases, with degradation in each of these phases appearing to follow first-order kinetics, a model commonly applied to describe virus inactivation (28). To accommodate the difference in inactivation rates, we used saliva droplet drying time as the breakpoint to distinguish kinetics in these two phases. Simple linear regression of the data prior to droplet drying indicates that influenza virus in saliva does not decay significantly during the wet phase (slope = 0.010 ± 0.012 min^-1^; mean ± 95% CI), but after drying, the inactivation rate increases to 0.036 ± 0.020 min^-1^.

In contrast to the trends observed in saliva droplets, decay in airway surface liquid continued at a similar rate between the two phases, also following first-order kinetics. We were not able to detect any differences in inactivation rates between the wet and dry phases for airway surface liquid, likely due to the low level of inactivation observed over exposure period of one hour. While decay of virus in airway surface liquid droplets was low, the virus inactivation rate was significantly greater than zero, 0.010 ± 0.0030 min^-1^. Our findings confirm the protective effect of airway surface liquid in comparison to saliva at intermediate RH. Additionally, they highlight considerable differences in the inactivation of influenza virus in saliva during the wet and dry phases and emphasize that the majority of decay at intermediate RH is driven by mechanisms that occur during the dry phase.

**Figure 4.**
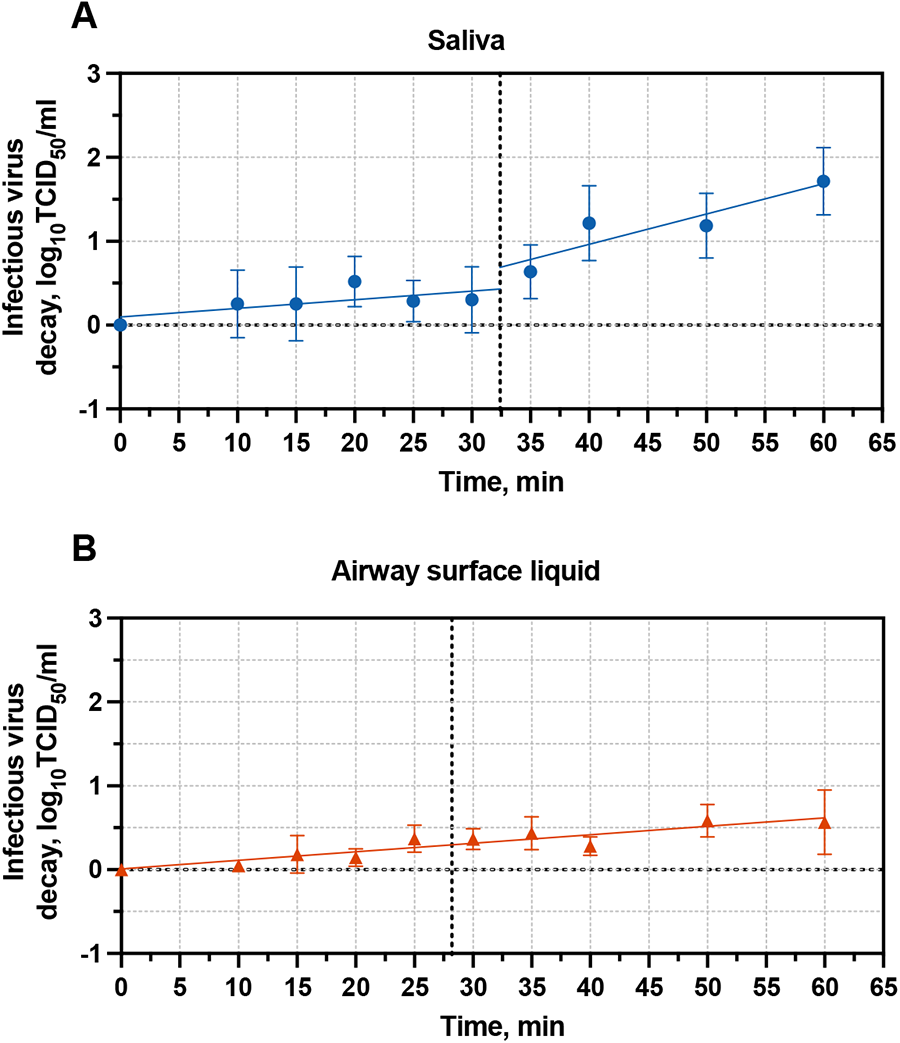
Inactivation kinetics of H1N1pdm09 at 50% RH in 1 µL droplets. Droplets were comprised of (A) human saliva or (B) airway surface liquid. Three technical replicates were conducted for each independent replicate, and the mean and standard deviation of five independent replicates are shown. Simple linear regression analysis was used to generate inactivation curves. Breakpoints in inactivation curves were established by average droplet drying time from two independent replicates (Table S1) and are shown as dotted vertical lines.

### Droplet composition and relative humidity influence droplet crystalline structure and nanobead location

It is possible the aggregation or interaction of viruses with other solutes in droplets upon drying could impact virus inactivation. Any observed differences in the crystal structure of airway surface liquid and saliva could therefore inform possible drivers of virus persistence. To assess how crystal structures within droplets vary by RH and droplet type, we captured microscopic images at 10x magnification of 1 µL droplets containing H1N1pdm09 comprised of human saliva or airway surface liquid following a two-hour exposure to 20% RH, 50% RH, and 80% RH environments (Fig. 5). In line with our evaporation measurements, droplets did not completely dry out at 80% RH, leading to a gelatinous solution without much discernible structure. Droplets at 20% RH and 50% RH, on the other hand, exhibited extensive crystalline structure. Dried airway surface liquid droplets had feather-like crystals once dry at 20% and 50% RH, while saliva displayed skinnier, line-like crystalline structures. These line-like crystalline structures in the saliva droplets differed at 20% and 50% RH, with more densely packed structures at 20% RH. In addition, a thick ring of structure was observed at the edge of airway surface liquid droplets at 20% and 50% RH, referred to as a “coffee-ring” effect (i.e., movement of solutes within the droplet to the liquid-surface edge) (29, 30), while a thin coffee-ring was present at the edge of saliva droplets at 20% and 50% RH.

**Figure 5.**
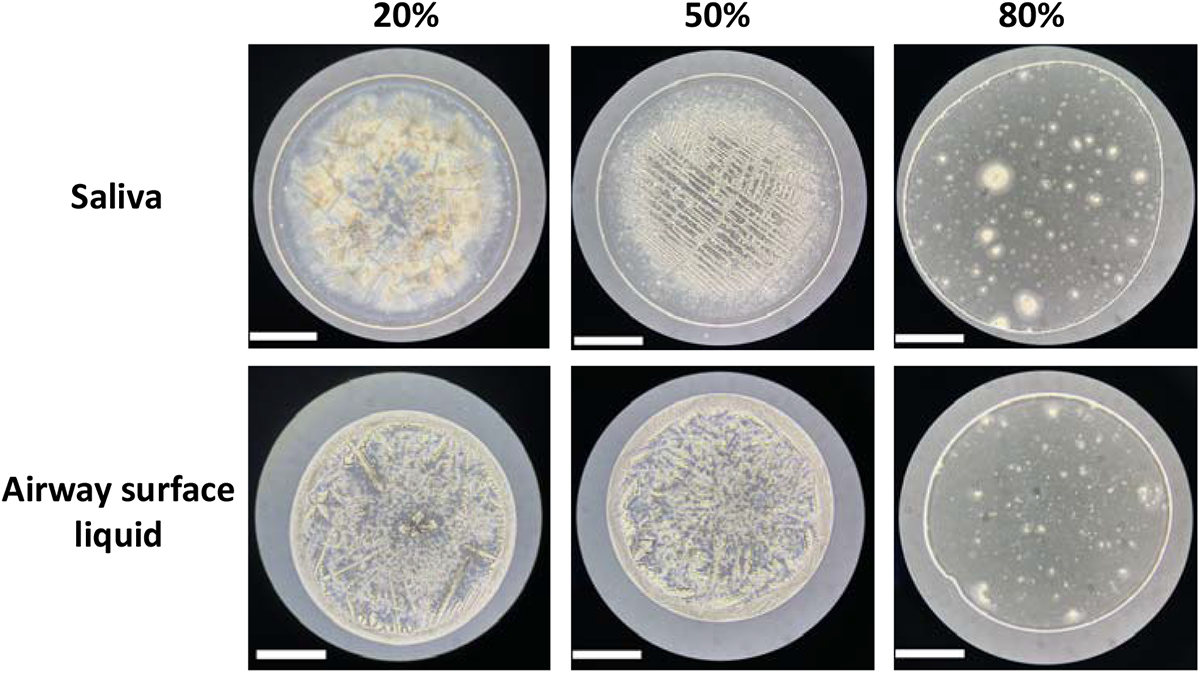
Bright field images of droplets comprised of human saliva or airway surface liquid containing H1N1pdm09 virus after exposure to 20%, 50%, and 80% RH for two hours. Images were taken at 10x magnification. Scale bars, 500 µm.

The differences in crystal structure formed at variable RH could play a role in virus persistence if virus interaction or incorporation with these crystal structures impacts inactivation. In situ virus particle visualization in droplets presents considerable challenges because of the small size of virions. We therefore used fluorescent nanobeads similar in size to influenza virions as a proxy for virus particles; these beads have a strong fluorescent signal that is easily detectable using standard fluorescence microscopy techniques. In both respiratory fluids, nanobeads clustered around the outer rim of the droplet upon drying, regardless of RH (Fig. 6, Fig. S3). At 80% RH, the coffee-ring distribution was more dispersed than at 20% RH and 50% RH. This is likely due to the fact that the droplets had not completely dried by observation, leading to a gelatinous solution lacking in crystalline structures that would allow for nanobead colocalization. At 20% RH and 50% RH, extensive nanobead accumulation in the airway surface liquid droplets was also observed in the interior of the droplet, where beads colocalized with crystalline structures. To a lesser extent, this phenomenon was also observed in saliva. While the nanobeads in the coffee-ring of the airway surface liquid droplets coincided with the thick crystalline complex located around the perimeter of the droplet, the nanobeads in saliva droplets did not appear to collocate with crystalline structures in the coffee-ring, but rather collocated with the thin region around the perimeter of the droplet that appeared to be free of any crystalline structures. Additional work to identify exactly which solutes or proteins are found in these regions of the different droplets will be beneficial to uncovering potential interactions of virions and particles that could play an important role in virus inactivation. Together, these data indicate that distinct respiratory fluids exhibit composition-dependent differences that likely influence influenza virus localization, interactions, and stability.

**Figure 6.**
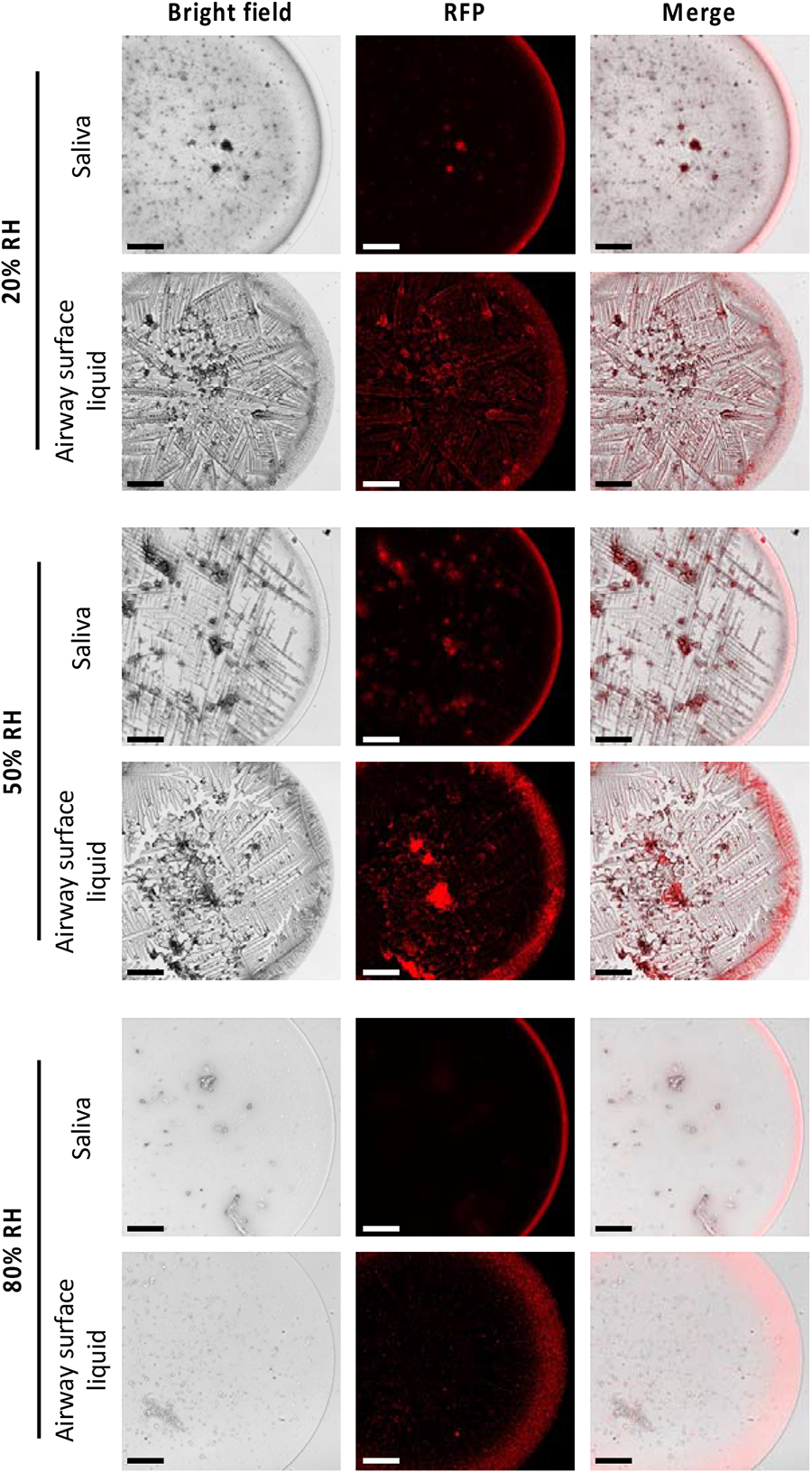
Fluorescence images of droplets comprised of human saliva or airway surface liquid with 100 nm fluorescent nanobeads (1:1000 dilution) after drying at 20%, 50%, or 80% RH for two hours. Images were taken at 10x magnification. RFP = red fluorescence protein. Scale bars, 200 µm.

## Discussion

Little work to date has looked at infectious influenza virus levels present in the oral versus nasal cavities of infected hosts over time, despite the fact that respiratory fluid particles composed of saliva could contribute to onward spread. Limited clinical work has identified influenza virus nucleic acid or antigen in oral specimens (31–33). Another study focused on culture-based detection of influenza in humans did consistently find nose and throat samples positive for infectious virus (34). Here, we consistently detected infectious virus in oral swabs, as opposed to oropharyngeal swabs, sampled from ferrets during influenza A virus infection. Our findings confirm that influenza virus present in the oral cavity of an infected host is likely to be infectious, lending experimental evidence for the possibility of onward transmission from both nasal and oral cavity expulsions. Future work focused on the quantity, size distribution, and composition of emissions from each cavity is needed to ascertain the relative contribution of each anatomical site to virus-laden particles in the environment.

In this study, we also compared the environmental persistence of influenza virus in droplets of saliva versus airway surface liquid, the latter being representative of mucus-containing respiratory fluid. We demonstrated that two human seasonal influenza A virus strains, H1N1pdm09 and H3N2, are more susceptible to decay at midrange humidity in saliva as compared to in airway surface liquid. Influenza virus degradation in airway surface liquid droplets was similar to the levels observed in previous studies from our laboratory (19). Recent work investigating bacteriophage persistence in saliva microdroplets (i.e., particles ranging from ∼ 30 to ∼ 600 µm in diameter) deposited on a glass surface demonstrated robust stability, with three different bacteriophages decaying by at most 1.5-log_10_ after 14 hours over a broad range of humidities (21). Phages exhibited more decay in a buffer solution and water compared to saliva at all RHs, the inverse of what we observed here. Our use of larger droplet sizes (∼ 1.3 mm), distinct viruses, and/or dissimilar saliva sources could explain the differences observed, although further investigation is required. On the other hand, work with vesicular stomatitis virus in 2 µL saliva droplets at variable RH resulted in the same U-shaped trend in virus decay that we observed with influenza virus in saliva (23). Murine coronavirus in saliva aerosols also exhibited reduced decay at 20% RH compared to at 50% or 80% RH (22). Taken together, these observations suggest there may be substantial differences in virus fate depending on respiratory fluid composition, RH, and droplet or aerosol size. Future work addressing these distinctions, specifically identifying the composition of different sized respiratory particles (35), is needed to accurately define transmission risks from viruses in the environment.

By more carefully examining the kinetics of droplet drying and virus decay in saliva and airway surface liquid, we found that while both fluids demonstrated similar drying times, there were distinct differences in influenza virus persistence patterns. At midrange humidity, virus in saliva droplets exhibited minimal inactivation during the wet phase, but upon drying, decay increased significantly. In contrast to virus in saliva, virus decay occurred at sustained, low levels in airway surface liquid regardless of droplet phase. From these data, we cannot be certain that virus in airway surface liquid does not exhibit a shift in inactivation kinetics following drying; rather the overall virus reductions were too low for us to dependably detect a change. Distinct inactivation trends in different phases of droplet drying have also been observed for influenza and other respiratory viruses in cell culture medium, although the trends are not consistent (13, 27). The decay rates we detected for influenza virus in airway surface liquid and saliva droplets during the dry phase were elevated compared to those from a recent study for influenza virus in cell culture medium (27), which could be due to different droplet composition and/or the timeframe measured. The differences we observed in virus inactivation between the wet and dry phase of saliva droplets underscore the importance of the virus’ microenvironment in understanding the mechanisms governing virus persistence. Additionally, our results highlight the need to study virus persistence in physiologically relevant solutions, as different matrices clearly exhibit altered trends in virus decay.

Many factors have been proposed to drive environmental virus decay in droplets or aerosols, but the true causes of inactivation remain unknown. Yang and Marr showed reduced degradation of influenza virus in droplets with added protein at intermediate RH compared to those without protein (15). Increased salt concentrations in aerosols have correlated to increased inactivation of Langat virus but protection of poliovirus and T7 bacteriophage (11). Our work provides additional insights; despite similar bulk protein levels across respiratory solutions, influenza virus decay was strikingly distinct between airway surface liquid and saliva droplets. In addition, the reduced salt concentrations in saliva as compared to airway surface liquid indicate salt content does not explain the inactivating effect on influenza virus here, though it is possible the differences in salt concentration from saliva to airway surface liquid were not sufficient to significantly affect virus persistence. Indeed, Benbough et al. investigated the impact of differences in salt content on the order of 50 ppt (11); saliva and airway surface liquid only had differences of ∼ 8 ppt. Our findings indicate bulk protein or salt content concentrations alone are unlikely to have driven our virus inactivation results.

The exact composition of chemicals and proteins present in saliva and airway surface liquid, however, could impact virus persistence. Specifically, the increased virus decay observed in saliva could be due to the presence or absence of constituents with protective or antiviral properties that differ from those in airway surface liquid. Previous work has described anti-influenza virus activity in saliva acting by hemagglutination inhibition and virus neutralization (36–38). This research identified specific antiviral salivary components, including protein-bound sialic acid, salivary scavenger receptor cysteine-rich protein (gp-340), and gel-forming mucin 5B (MUC5B), that display antiviral properties (36, 37, 39). A recent study focused on vesicular stomatitis virus decay in saliva provided evidence for extensive antiviral protein-mediated decay in droplets at intermediate RH that may be partially associated with lysozyme activity (23). The protective and antiviral factors comprising airway surface liquid have not been as well-studied, although proteomic analysis of airway surface liquid identified several dominant mucins, with MUC5B being the most abundant (40). A comparative analysis of proteins in distinct respiratory fluids would prove useful in characterizing constituents in these solutions with protective or antiviral activity that may impact environmental virus persistence.

How these specific antiviral components of saliva play a role in the distinct patterns of influenza virus decay observed in droplets at various RHs is currently unclear. Their potential presence does not explain the reduced decay measured in saliva at 20% RH in our study, unless more rapid evaporation of droplets at low RH resulted in reduced activity or interactions with virus prior to and after dehydration. Interestingly, our controls of virus suspended in bulk saliva solution at ambient temperature resulted in little to no reduction in infectious virus over the experimental period (Fig. S4). This suggests that the concentrations of antiviral constituents in bulk saliva were not sufficient alone to extensively reduce influenza virus infectivity during this time period and that RH-mediated evaporation changes in small droplet volumes contribute significantly to decay.

The different crystalline structures we observed in droplets and the corresponding virus associations with those structures (or with other factors within them) could impact virus inactivation in the dry phase. Indeed, we noted that the morphology of dried droplets and distribution of 100 nm nanobeads, a proxy for influenza virus particles, differed considerably across RHs and droplet compositions. Huang et al. previously described a relationship between the coffee-ring effect and virus persistence, demonstrating that for different media-based droplet solutions, a thicker coffee-ring upon drying was correlated with less overall decay of the commonly used surrogate virus bacteriophage phi6 (29). In contrast, when spiking vesicular stomatitis virus and lysozyme, an antiviral protein, into saliva droplets, Kong et al. observed increased colocalization of virus and lysozyme in the coffee-ring of evaporated saliva droplets at 50% RH compared to 20% RH; this was hypothesized to explain the increased virus decay observed at intermediate RH (23). While we observed the thickest coffee-ring was associated with dried airway surface liquid droplets and their viral persistence, virus-containing saliva droplets at 20% RH exhibited a minimal ring but viruses retained infectivity. Additionally, the spatial distribution of our fluorescent nanobeads generally followed the coffee-ring effect, although nanobeads were observed throughout the airway surface liquid droplets in complexes that could be proteinaceous or solute-derived, particularly at 50% RH. Clearly the significance of the coffee-ring of solutes may vary depending on the matrix composition and virus used. Furthermore, not all droplets undergo the coffee-ring effect upon drying, and perhaps these differences in particle colocalization explain the differences in influenza inactivation observed in airway surface liquid and saliva droplets at variable RH. It is important to note that the colocalization observed in this study and in other work does not confirm interactions of these constituents in dried droplets. Future work characterizing the interactions between virions and antiviral or protective constituents in distinct respiratory fluid droplet matrices will be critical to establishing the mechanisms driving inactivation during the droplet’s dry phase.

While we have established important differences in influenza virus persistence in droplets comprised of distinct respiratory fluids, our work has limitations. We studied influenza virus persistence in 1 µL droplets, which are large (i.e., ∼ 1240 µm in diameter) compared to the range of aerosol and droplet sizes that has been measured during respiratory activities (4). Past work with influenza virus has shown similar stability trends in aerosols and droplets (19), however we cannot be certain the results we have observed in 1 µL droplets will hold in smaller droplets or aerosols. Future research efforts should focus on representative droplet and aerosol particle sizes using relevant respiratory fluids, like those used in this work, to ensure our findings are generalizable across a range of particle sizes. In addition, while our work holds for the two seasonal influenza A viruses assessed, persistence in respiratory fluids should be assessed for other strains and additional important respiratory viruses, particularly for those with distinct genome types and structures, including rhinoviruses, adenoviruses, and coronaviruses.

Environmental persistence of influenza viruses is hypothesized as one possible factor contributing to sustained virus transmission in temperate regions during winter months (41, 42). Our findings could help explain these seasonal trends. If onward transmission of influenza is driven by aerosols and droplets predominantly comprised of saliva, then the decreased persistence of influenza at midrange humidity would be in line with reduced transmission observed in summer months, when indoor RH typically ranges from 40% to 60% RH (41). Increased stability of influenza virus in saliva droplets at low RH would be consistent with sustained transmission in winter months, when indoor RH is usually between 10% and 40% (41). While our findings of influenza persistence in saliva align with observed seasonal trends, the ubiquitous survival of influenza virus in airway surface liquid does not. Additional research characterizing the relative proportion of different respiratory fluids in virus-laden aerosols or droplets contributing to onward transmission is needed to understand whether environmental persistence plays a significant role in influenza virus seasonality. In addition, characterization of the aerosol size distribution expelled by the mouth versus the nose would help inform the use of correct aerosol sizes and provide insight into the relative composition of aerosols from these cavities. This work has important implications for developing robust interventions to mitigate transmission during peak periods of influenza virus illness and spread.

## Materials and Methods

### Influenza virus stocks and quantification

Biological influenza virus A/California/07/2009 (H1N1pdm09) and recombinant influenza virus A/Perth/16/2009 (H3N2) were used in this study as previously described (20). Virus propagation was conducted by growing 1:50,000 CP1 virus stocks on confluent Madin-Darby canine kidney (MDCK) cells, kindly provided by Dr. Kanta Subbarao, in infection medium (Eagles’ Minimum Essential Medium (MEM) with L-glutamine (Lonza, Cat. No. BE12-611F) containing 2x antibiotic-antimycotic (Gibco, Cat. No. 15240062), L-glutamine (Lonza, Cat. No. BE17-605E), and TPCK trypsin (Worthington-Biochem, Cat. No. LS003750)) at 37°C. Virus was harvested after significant cytopathic effect was observed. Cellular material was removed through centrifugation at 2,000 x g for 10 minutes at 4°C. The H3N2 virus stock was concentrated through a 30% sucrose cushion to increase the detectable range of virus decay in droplet experiments by ultracentrifugation at 24,000 x rpm for 1.5 hours at 4°C after propagated virus was harvested from cells. Following ultracentrifugation, H3N2 virus was resuspended in infection medium and vortexed vigorously. Virus stocks were stored in aliquots at -80°C until use. Infectious virus concentrations were quantified using the tissue culture infectious dose 50 (TCID_50_) Spearman Karber method, as previously described (43).

### Animal ethics statement

Ferret experiments were conducted in a biosafety level 2 facility at the University of Pittsburgh in compliance with the guidelines of the Institutional Animal Care and Use Committee (approved protocol 22061230). Animals were sedated with isoflurane following approved methods for all nasal washing and oral swabbing.

### Animal work

Ferrets were infected intranasally with ∼ 10^6^ TCID_50_/500 µL of influenza virus (H3N2 or H1N1pdm09), 250 µL administered into each nostril. Ferret nasal cavities and oral cavities were sampled for influenza virus on days 1, 2, 3, 5, and 7 post-inoculation. Specifically, ferrets were anesthetized with isoflurane, and Floq Swabs (Copan, Cat. No. 525CS01) were used to swab each infected ferret’s hard and soft palate, cheeks, and tongue. Swabs were immediately placed in 500 µL Eagles’ Minimum Essential Medium (MEM) with L-glutamine (Lonza, Cat. No. BE12-611F) containing 2x antibiotic-antimycotic and L-glutamine. Following oral swab collection, nasal washes were conducted by collecting 1 mL Dulbecco’s phosphate-buffered saline (DPBS; Gibco, Cat. No. 14190250) through the ferret’s nasal cavity and collecting flushed nasal solution. Oral swab and nasal wash samples were stored at -80°C until TCID_50_ analysis.

### Bulk respiratory fluid analyses

The total protein concentration in each respiratory solution was determined by the bicinchoninic acid assay (BCA; Thermo Scientific, Cat. No. 23225), using bovine serum albumin (Thermo Scientific, Cat. No. 23210) as the protein standard. Conductivity, total dissolved salt, salinity, and pH were quantified with a field probe (ThermoFisher, Cat. No. 13-643-124).

### Droplet generation

Pooled human saliva (Innovative Research, Cat. No. IR100044P) or airway surface liquid (details of respiratory fluids provided below) was used for virus droplet solutions. In each independent experiment, virus stocks were diluted 1:10 into each droplet solution. 10 x 1 µL droplets of each diluted suspension were deposited on polystyrene material (6-well plates, Thermo Scientific, Cat. No. 140675) and exposed to ambient temperature and variable RH (20%, 50%, or 80%) for one hour or two hours in a controlled environmental chamber (Electro-Tech Systems, 5532 Series). A Hobo Temperature and Humidity Data Logger (Onset, Cat. No. UX100-011) was used to monitor the temperature and RH for the duration of each experiment (Fig. S2). After the exposure period, droplets were resuspended in 500 µL of MEM with L-glutamine and immediately stored at -80°C until analysis. Droplets were also generated and collected at zero hours. Three independent replicates were completed with technical triplicates conducted for each RH condition and droplet composition. Control samples of the bulk droplet solution were collected at zero and two hours to ensure no significant influenza virus occurred.

### Respiratory fluids

Pooled human saliva was processed as specified by Innovative Research. Briefly, saliva samples were stored at -80°C upon collection and pooled by thawing samples and combining them. The pooled solution was then passed through a cheese cloth before aliquoting and freezing at -80°C until purchase. Details regarding the number of samples and information about saliva donors were not disclosed.

Human bronchial epithelial (HBE) cultures were grown at the air-liquid interface as previously described (19). Airway surface liquid was collected periodically from the air-liquid interface by adding 100 – 150 µL phosphate buffered saline to wells, incubating cells for 5 min at 37°C, and harvesting the solution following incubation. Airway surface liquid was collected from HBE cultures from multiple donor patients to capture potential heterogeneity in viral persistence. Specifically, airway surface liquid droplet suspensions used in each independent replicates were from a different donor HBE culture. Five different HBE cultures were used in influenza virus persistence work (deidentified culture numbers: 0277, 0284, 0302, 0304, and 0305), and each independent replicate was performed in triplicate.

### Examination of droplet drying kinetics

10 x 1 µL droplets comprised of virus solutions, generated as described above using H1N1pdm09 virus in either airway surface liquid or saliva, were deposited on a 35 mm x 10 mm polystyrene petri dish (Falcon, Cat. No. 351008) and placed on a microbalance (Sartorius Cubis I, Model MSE3.6P, readability = 0.0010 mg) within the environmental chamber. The chamber was set at 20%, 50%, or 80% RH. The glass enclosure of the microbalance was removed to ensure droplets were exposed to the desired RH within the chamber. Droplet mass was recorded every 10 seconds for two hours. The time to droplet drying was determined by assessing the point at which the droplet mass left the linear phase of evaporation. Two independent replicates were conducted for each RH condition and droplet composition. Images of the droplets were taken every 10 seconds to visually observe changes in droplet morphology and drying and are available in supplemental videos S1 – S6. While the microbalance was zeroed prior to each experiment, balance sensitivity immediately after droplet addition often resulted in droplet masses dropping below zero. In these cases, the minimum mass achieved over the two-hour experimental period was assumed to be the zeroed mass for calculations of percentage original mass. This had no impact on the drying time, which was only dependent on when the rate of mass loss diverged from the droplet’s linear evaporation phase. The percentage of original mass, shown in Fig. 3, was determined using the following equation:

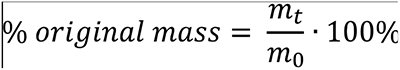

Where the initial droplet mass and droplet mass at time t were m_0_ and m_t_, respectively.

### Short-term inactivation kinetics

10 x 1 µL droplets comprised of H1N1pdm09 diluted 1:10 in human saliva or airway surface liquid were deposited on 6-well cell culture plates. Technical triplicates were carried out for each of five independent replicates. Droplets were collected at 0, 10, 15, 20, 25, 30, 35, 40, 50, and 60 mins following exposure to 50% RH in the environmental chamber. RH and temperature were logged with a Hobo Temperature and Data Logger. One droplet exposure time was conducted at a time, and sample exposure times were randomized. First-order decay was used to fit the data by the following equation:

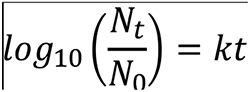

Where N_0_ and N_t_ are the influenza virus concentrations in droplets at time zero and time t, respectively, and k represents the inactivation rate constant.

### RNA recovery in droplets after drying

Experiments were conducted before and after droplet drying to ensure droplet recovery was the same regardless of droplet desiccation state. 10 x 1 µL droplets containing H1N1pdm09 diluted 1:10 in MEM or respiratory fluid were deposited on 6-well tissue culture plates as described above. Droplets were collected following exposure to 50% RH for zero and one hour. Three independent replicates were conducted with each droplet type. Droplets were collected in 500 µL MEM and stored at -80°C until extraction. RNA extractions were carried out with the QIAamp Viral RNA Mini Kit (Qiagen, Cat. No. 52904) following the standard extraction protocol. Extracts were eluted in 60 µL nuclease-free water and stored at -80°C until RT-qPCR analysis. RT-qPCR targeting the matrix gene (primer and probe sequences in Table S2; modified from previously published primers/probe (44)) was performed using the BioRad iTaq Universal Probes One-Step Kit (BioRad, Cat. No. 1725140). Thermocycling was performed on a BioRad CFX Connect Real-Time PCR Detection System (BioRad, Cat. No. 1855201) with the following cycle settings: reverse transcription at 50°C for 10 min, initial denaturation at 95°C for 2 min, 40 cycles of denaturation and annealing/extension at 95°C for 10 sec and at 60°C for 20 sec, respectively. In vitro transcribed RNA of the full-length matrix gene was used as the standard.

### Bright field microscopy of dried droplet structure

To visualize droplet morphology after drying, 1 µL droplets comprised of virus solutions, generated as described above using H1N1pdm09 virus in either airway surface liquid or saliva, were deposited on polystyrene chamber slides (Ibidi, Cat. No. 80826) and exposed to 20%, 50%, or 80% RH for two hours at ambient temperature. Droplets were immediately visualized using an inverted microscope (Olympus, Model CKX53) at 10x magnification.

### Nanobead visualization in dried droplets

100 nm carboxylate-modified, fluorescent, polystyrene nanobeads (Invitrogen, Cat. No. F8801) were sonicated for 10 min and subsequently spiked into airway surface liquid or saliva at a final 1:1000 dilution. 1 µL droplets from these solutions were then generated on polystyrene chamber slides. Droplets were immediately placed in the environmental chamber and exposed to 20%, 50%, or 80% RH for two hours. The chamber was kept dark during the exposure. Following the two-hour exposure, fluorescent nanobeads in droplets were visualized using an inverted fluorescence microscope (Olympus, Model IX73P2F) at 10x magnification. Saliva and airway surface liquid droplets without nanobeads were used as negative controls.

### Statistical analyses

All statistical analyses were conducted in Prism Version 9.2.0.

## Acknowledgments

This work was funded in part with Federal funds from the National Institute of Infectious Diseases, National Institutes of Health, Department of Health and Human Services, under Contract No. 75N93021C0015 and the FluLab, Mitigate Flu Project. We thank Dr. Rachel Duron and Lakdawala laboratory members for critical review and feedback.

